# Nucleolar Stress-inducing Compounds Influence rDNA occupancy of RNA Polymerase I Transcription Machinery

**DOI:** 10.1101/2025.01.09.632225

**Authors:** Matthew V. Yglesias, Victoria J. DeRose

## Abstract

Transcription of ribosomal RNA (rRNA) by RNA Polymerase I (Pol I) is often upregulated in cancer to facilitate rapid cell growth and proliferation, and has emerged as a potential target for chemotherapeutic agents. BMH-21 and Pt(II) chemotherapeutic agent oxaliplatin are well documented as inhibitors of Pol I activity, however the underlying mechanisms for this inhibition are not completely understood. Here, we applied chromatin immunoprecipitation sequencing (ChIP-seq) techniques and immunofluorescence imaging to probe the influence of oxaliplatin and BMH-21 on Pol I machinery. We demonstrate oxaliplatin and BMH-21 induce early nucleolar stress leading to the formation of “nucleolar caps” containing Pol I and upstream binding factor (UBF) which corresponds with broad reductions in ribosomal DNA (rDNA) occupancy of Pol I. Distinct occupancy patterns for the two compounds are revealed in ChIP-seq experiments. Taken together, our findings suggest that in vivo, oxaliplatin does not induce Pol I inhibition via interrupting a specific step in Pol I transcription, while treatment with BMH-21 induced unique polymerase stalling at the promoter and terminator regions of the human ribosomal RNA gene.

## Introduction

Ribosome biogenesis is a highly coordinated essential process which involves the synthesis, processing and assembly of numerous ribosomal RNAs (rRNA) and proteins into functional ribosomes. The majority of rRNA is transcribed by RNA polymerase I (Pol I), which accounts for nearly 50% of all RNA transcription in the cell (Russell & Zomerdijk, 2005). As the initial and rate-limiting step of ribosome biogenesis, the rate of rRNA transcription is a key mediator in ribosome production, and is proportional to cell growth and proliferation (Lafontaine et al., 2021; Pitts & Laiho, 2022).

In humans Pol I solely transcribes the 47S pre-rRNA from ribosomal DNA genes (rDNA), which are organized in clusters of tandem repeats situated on the short arm of acrocentric chromosomes (Xuan et al., 2021). The 47S pre-rRNA undergoes several co– and post-translational processing steps to generate the mature 18S, 5.8S, and 28S rRNAs; the 5S is transcribed separately by RNA polymerase III (Penzo et al., 2019). Pol I transcription takes place in the nucleolus, a membrane-less organelle within the nucleus, which serves as the primary site of ribosome biogenesis. The nucleolus is organized into three nested layers maintained in part via liquid-liquid phase separation (LLPS): the fibrillar center (FC), dense fibrillar component (DFC) and the granular component (GC) (Lafontaine et al., 2021). The organization of nucleolar subcomponents reflects the individual steps in ribosome biogenesis. Initial Pol I transcription of pre-rRNA takes place in the FC, near the border of the DFC, to allow for nascent pre-rRNA to be co-transcriptionally processed in the DFC (Pitts & Laiho, 2022). Mature rRNAs migrate into the GC where they, along with the 5S rRNA, are assembled with ribonucleoproteins to generate the small 40S and large 60S pre-ribosomal subunits. The pre-40S and pre-60S subunits then enter the nucleoplasm, where they undergo late-stage maturation before final export into the cytoplasm where they combine to form fully functional ribosomes (Penzo et al., 2019).

To facilitate the increased translational and metabolic demands of tumorigenesis, cancer cells often display dysregulations in ribosome biogenesis and heightened rates of rRNA synthesis, which also functions as an important clinical biomarker (Penzo et al., 2019). The inherent instability of rDNA, coupled with the hyperactivation of ribosome biogenesis, make cancer cells particularly susceptible to disruptions in ribosome biogenesis and Pol I activity, which has found growing interest as a potential chemotherapeutics target (Hwang & Denicourt, 2024; Xuan et al., 2021).

Several small molecule and clinically relevant drugs have been reported to target Pol I and rRNA synthesis. Actinomycin D (ActD), widely used as an RNA transcription inhibitor, blocks polymerase transcription elongation by intercalating into DNA. RNA Pol I is ∼10× more sensitive to ActD compared to RNA Pol II and III, which effectively allows for selective inhibition of Pol I when treating at low concentrations of ActD (Burger et al., 2010; Bensaude, 2011).

BMH-21, a quinazolinone derived DNA intercalator, has been shown to inhibit rRNA transcription by disrupting Pol I activity, leading to loss of rDNA occupancy and subsequent degradation of the RNA Pol I subunit, RPA194 (Jacobs, Huffines, et al., 2022; Peltonen et al., 2014; Wei et al., 2018). *In vitro* studies further demonstrated that BMH-21 selectively inhibits Pol I transcription initiation, promoter escape, and elongation (Jacobs, Fuller, et al., 2022; Jacobs, Huffines, et al., 2022).

The small molecule intercalator CX-5461 was identified in a screen for inhibitors of rRNA transcription and was initially characterized as a selective Pol I inhibitor. CX-5461 stabilizes G-quadruplexes found in rDNA, and disrupts the Pol I initiation complex (PIC) by preventing promoter binding and release (Drygin et al., 2009, 2011; Jean-Clément Mars et al., 2020). In additional studies, the cytotoxic effects of CX-5461 have been attributed to topoisomerase II (TOP2) poisoning (Bruno et al., 2020), and implicated in the inhibition of TOP2*α* associated with Pol I in the mechanism of action (Cameron et al., 2024).

Pt(II)-based chemotherapeutics have long been known to cause disruptions in ribosome biogenesis and rRNA synthesis, at elevated concentrations (Burger et al., 2010; Jordan & Carmo-Fonseca, 1998; Peterson et al., 2015). However, this was not considered a primary mechanism for cytotoxicity until later work identified oxaliplatin as inducing cell death specifically through a disruption in ribosome biogenesis, in contrast to cisplatin and other Pt(II) chemotherapeutics which act by triggering the DNA damage response (DDR) (Bruno et al., 2017). Later work reinforced these findings, demonstrating that oxaliplatin induces nucleolar stress—a hallmark for disruption of ribosome biogenesis— that proceeds from the rapid inhibition of nascent rRNA synthesis (Sutton et al., 2019; Sutton & DeRose, 2021).

Some studies suggest that oxaliplatin-induced inhibition of rRNA synthesis is not due to a specific molecular disruption of Pol I transcription, but rather caused by biophysical disruptions of nucleolar structure or function. Oxaliplatin has been shown to alter phase separation and dynamics of DFC and GC components leading to a disintegration of nucleolar subcomponents, which are able to disrupt Pol I activity and leading to activation of cell cycle arrest and apoptosis (Schmidt et al., 2022). However, alternative models have proposed that oxaliplatin inhibits Pol I transcription though activation of nucleolar DDR pathways involving the DDR kinase ATM/ATR signaling despite an absence of nucleolar specific DNA damage (Nechay et al., 2023). Others have demonstrated that rRNA transcription inhibition and nucleolar stress induction by oxaliplatin is not dependent on ATM/ATR activity (Pigg et al., 2024). Interestingly, DDR activation of ATM/ATR signaling has also been implicated in Pol I inhibition by CX-5461 (Cameron et al., 2024; Negi & Brown, 2015; Quin et al., 2016). Nucleolar activation of DDR and the resulting downstream effects are not well understood.

ActD, CX-5461, and BMH-21, show high affinity for GC-rich regions of the genome, such as in rDNA, which is purported to drive their selectivity and sensitivity for Pol I transcription (Bensaude, 2011; Goodisman et al., 1992; Peltonen et al., 2014). Pt(II) complexes also show selectivity for GC-rich regions of the genome but differ from DNA intercalators in their ability to form multiple covalent adducts with DNA through formation of 1,2-intrastrand crosslinks between adjacent purine nucleobases (Riddell & Lippard, 2018; Shu et al., 2016; Woynarowski et al., 1998).

The overall cellular mechanisms which dictate the various stress pathways induced by oxaliplatin, cisplatin, or other Pt compounds are not well understood, and a sufficient molecular understanding of rRNA transcription inhibition induced by oxaliplatin is currently lacking. Therefore, to better understand on how oxaliplatin perturbs Pol I activity we utilized chromatin immunoprecipitation sequencing (ChIP-seq) techniques to directly map the engagement or “occupancy” of rDNA transcription machinery along rDNA (Jean-Clement Mars et al., 2018; Sullivan & Santos, 2020), in comparison with BMH-21 as a well-characterized small molecule nucleolar stress inducer (Colis et al., 2014). In addition to mapping rDNA occupancy of the Pol I machinery, we utilized immunofluorescence imaging to characterize the connection between nucleolar function and morphology under treatment with Pol I inhibitors oxaliplatin and BMH-21.

High-throughput sequencing techniques such as native elongating transcript sequencing (NET-seq) have been adapted in *S. cerevisiae* for mapping Pol I occupancy on rDNA (Clarke et al., 2018), and have been previously used to elucidate the mechanism of Pol I transcription factor Spt4 (Huffines et al., 2021) and track the transcription rates of Pol I mutants (Huffines et al., 2022). NET-seq experiments performed on *S. cerevisiae* treated with BMH-21 revealed an acute reduction in Pol I occupancy as well as sequence-specific stalling of the Pol I elongation complex upstream of G-rich rDNA sequences (Jacobs, Huffines, et al., 2022). NET-seq is based on sequencing RNA, and reports on sites of active transcription. This method allows for precise probing of active Pol I occupancy at single-nucleotide resolution, but high levels of mature rRNAs have limited *in vivo* applications to yeast models expressing tagged-Pol I complex. Additionally, NET-seq may not capture changes in Pol I occupancy caused by Pol I inhibitors prior to transcription initiation. By contrast, ChIP-seq provides complementary information by detecting all DNA occupancy, but does not report on function of that occupancy.

In this study, ChIP analysis was performed using antibodies targeting proteins essential to rDNA transcription, Pol I subunit A (RPA194) and upstream binding factor (UBF). RPA194 is the largest catalytic subunit of the Pol I complex and is not found in RNA polymerases II or III (Pitts & Laiho, 2022). UBF is a member of the HMG-box DNA-binding protein family and is essential for mediating recruitment of the Pol I initiation complex to the rDNA promoter (Hamdane et al., 2014). UBF binds along the full rRNA gene, which maintains rDNA clusters in an open chromatin state to promote Pol I transcription, (Pitts & Laiho, 2022) and is purported to mark actively transcribing rDNA genes (Sanij et al., 2008).

Despite the fundamental importance of rRNA transcription and accessibility of sequencing techniques, relatively few ChIP-seq analyses of Pol I transcription machinery have been reported in the literature. ChIP-seq occupancy assays have been applied to mechanistic studies of CX-5461, and demonstrated that CX-5461 disrupts the Pol I transcription imitation complex by irreversibly blocking release of the promoter RNA polymerase I transcription factor (RRN3) (Jean-Clément Mars et al., 2020). ChIP-based occupancy assays of Pol I and related transcription machinery were also used to identify the transcription factor C/EAP alpha (CEBPA) as a factor in Pol I-RRN3 recruitment to rDNA (Antony et al., 2022).

Our findings suggest that BMH-21 and oxaliplatin disrupt rRNA synthesis through distinct mechanisms, which lead to the disengagement from rDNA of RNA Pol I, but not UBF. These effects were specific to ribosome biogenesis inhibitors and not observed in cisplatin treatments. This information may reveal a possible mechanistic target or pathway that may be beneficial to the development of new RNA Pol 1 specific reagents, or improve the efficacy of nucleolus-targeting therapeutics.

## Results and Discussion

### BMH-21 and oxaliplatin induce reorganization of FC components

Disruptions in ribosome biogenesis induce nucleolar stress and cause a redistribution of nucleolar subcomponents. A well-known hallmark for nucleolar stress is the relocalization of the nucleolar protein nucleophosmin (NPM1) from the GC to the nucleoplasm (Yang et al., 2018). Prior work found that an inhibition in rRNA synthesis by oxaliplatin corresponds with the induction of nucleolar stress, when utilizing the relocalization of NPM1 as a measure for quantifying nucleolar stress (Sutton & DeRose, 2021). To determine if relocalization of FC components follows a similar trend, we utilized immunofluorescence imaging to track the nucleolar localization of UBF and RPA194 in U2OS cells following treatment with 10 µM of cisplatin or oxaliplatin, or 1 µM of BMH-21.

Upon treatment with oxaliplatin or BMH-21, UBF and RPA194 condensed into round cap-like, puncta which localized at the periphery of the nucleolus, revealing a clear disruption in FC separation which was largely absent in cisplatin treatments at early timepoints. Commonly referred to as “nucleolar caps”, these structures are reported to form following inhibition of rRNA transcription or sustained rDNA damage (Boukoura & Larsen, 2024; Hwang & Denicourt, 2024; Lafontaine et al., 2021).

We observed initial formation of nucleolar caps in UBF-labeled images following 90 min treatment with BMH-21 and 3 h treatments with oxaliplatin (**Figure 1A**). Characteristic nucleolar caps were not observed in cisplatin treatments, which displayed a moderate condensation of UBF following 24 h treatment (**Figure 2A**).

**Figure 1.**
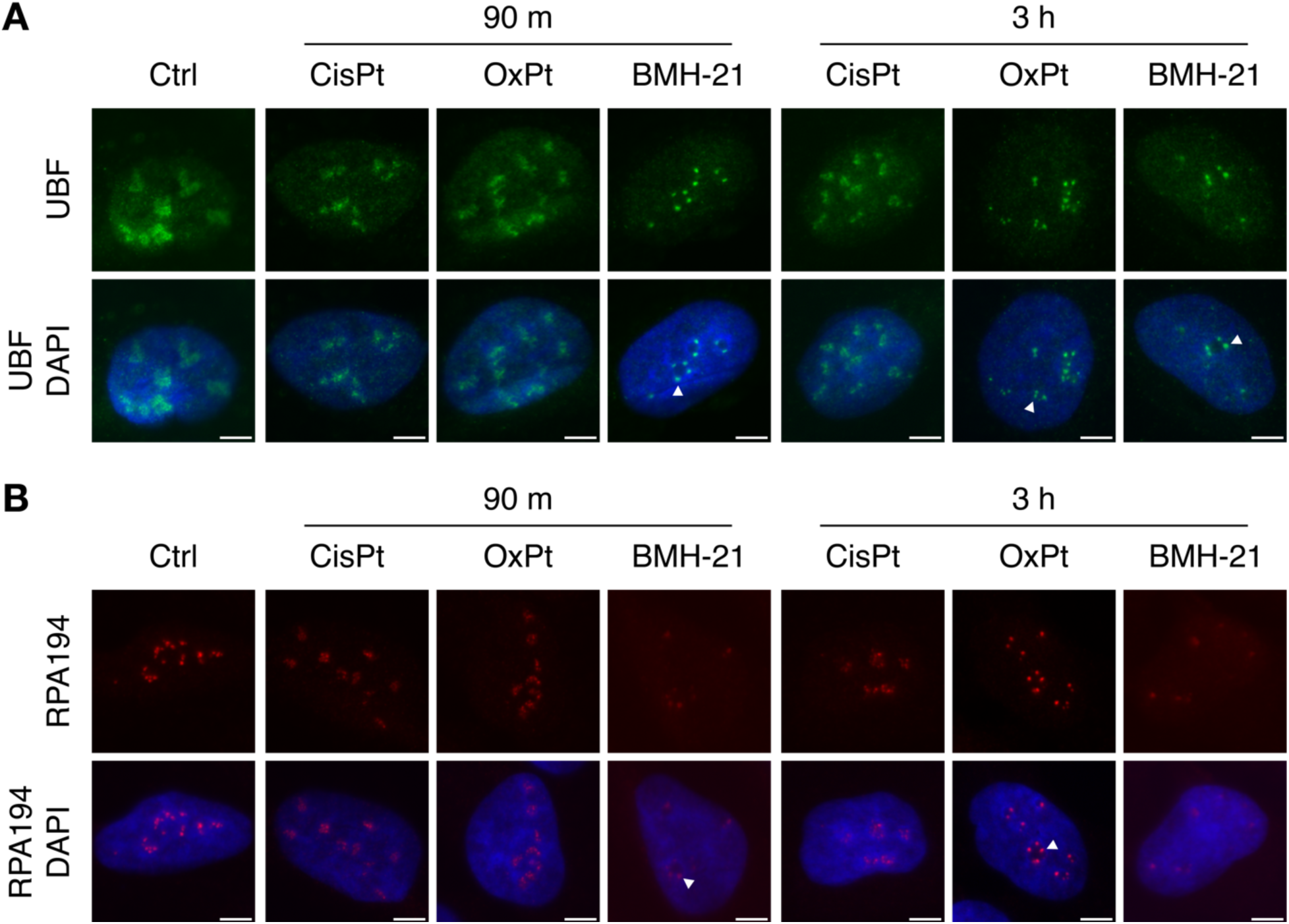
Oxaliplatin and BMH-21 induce early formation of nucleolar caps containing UBF and Pol I (RPA194). Representative U2OS cell images following 90 m and 3 h treatments with cisplatin (CisPt, 10 µM), oxaliplatin (OxPt, 10 µM), or BMH-21 (1 µM). Cells were immunostained for **(A)** UBF (green) or **(B)** RPA194 (red), and DNA (DAPI, blue). White arrow indicates nucleolar cap. Scale bars = 5 µm.

**Figure 2.**
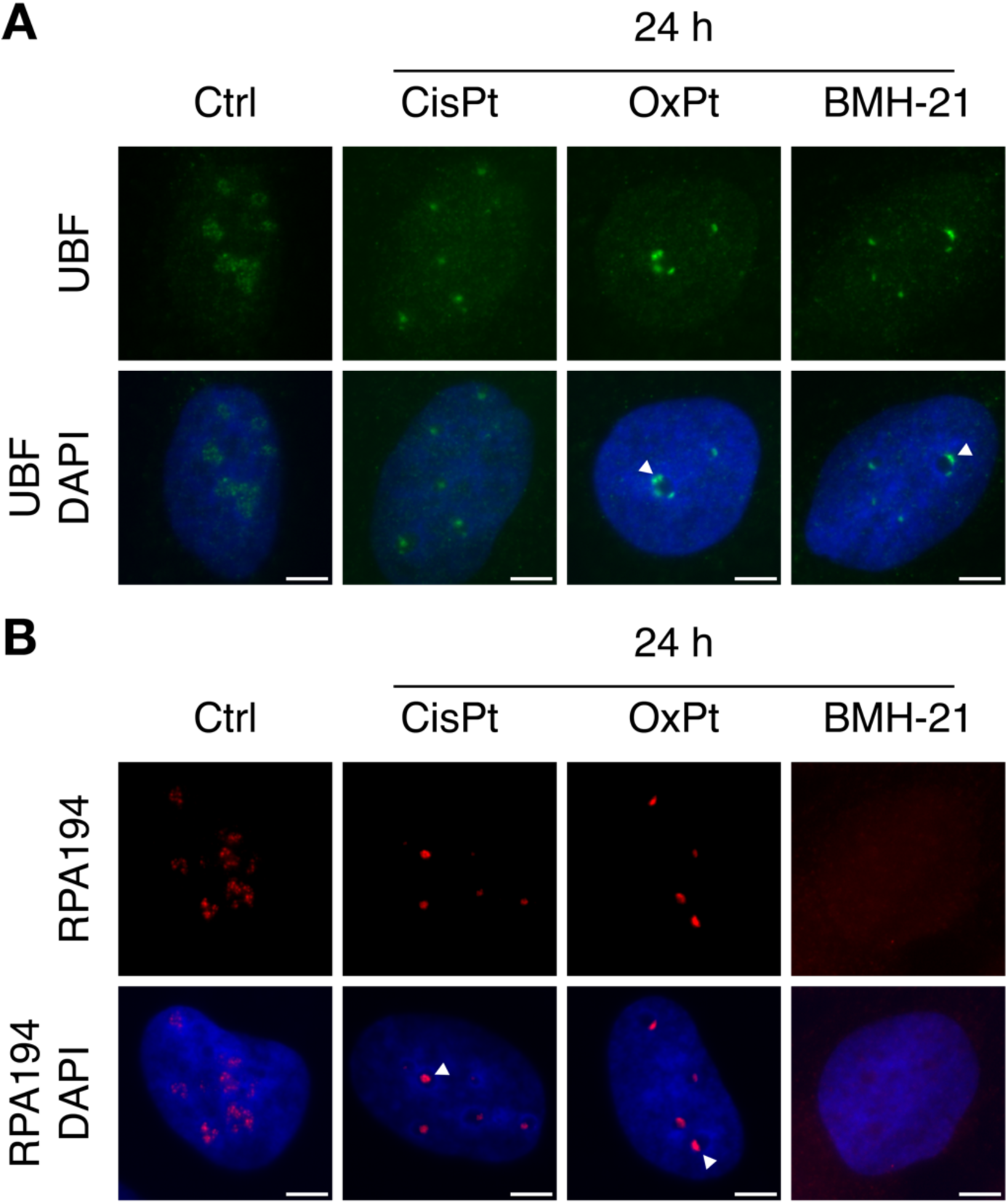
Representative images of UBF and Pol I (RPA194) localization in U2OS cells following 24 h treatment with cisplatin (CisPt, 10 µM), oxaliplatin (OxPt, 10 µM), or BMH-21 (1 µM). Cells were immunostained for **(A)** UBF (green) or **(B)** RPA194 (red), and DNA (DAPI, blue). White arrow indicates nucleolar cap. Scale bars = 5 µm.

The onset of nucleolar redistribution and cap formation was mirrored in cell images immunostained for RPA194. RPA194-containing nucleolar caps were initially observed following 90 min treatments with BMH-21 (**Figure 2B**). In agreement with previous reports demonstrating that a depletion of RPA194 coincided with formation of nucleolar caps with BMH-21 treatments (Colis et al., 2014; Wei et al., 2018), we also observed a loss in RPA194 immunofluorescence signal intensity, which was initially observed at 90 min timepoints (**Figure 1B**).

At 3 h timepoints, we found that oxaliplatin induced initial formation of nucleolar caps containing RPA194, while similar cap-like structures were only observed after 24 h treatment with cisplatin (**Figure 2B**). This later induction of nucleolar stress by cisplatin is consistent with our previous observations of nucleolar rearrangement occurring via downstream responses to DNA damage (Sutton et al., 2019; Sutton & DeRose, 2021). More robust RPA194-containing nucleolar caps were observed at later timepoints in oxaliplatin treatments as initial cap-like structures appeared to coalesce. This kinetic component of nucleolar reorganization in the nucleolar stress response has been previously observed in nucleolar stress induced by Pt(II) compounds (McDevitt et al., 2019; Pigg et al., 2022; Sutton & DeRose, 2021), where early induction of nucleolar stress was identified by relocalization of NPM1 in cells treated with oxaliplatin, but not cisplatin. The observations reported here indicate that oxaliplatin induces initial formation of nucleolar caps containing FC components UBF and RPA194 in a process that occurs at comparable timepoints to relocalization of GC component, NPM1.

### ChIP-based occupancy assay of Pol I transcription of rDNA

To directly investigate how rRNA transcription is inhibited by oxaliplatin we performed a ChIP-seq analysis on A549 and U2OS cells to map the rDNA occupancy of the Pol I machinery during early induction of nucleolar stress (Jean-Clement Mars et al., 2018). Based on previous work (Pigg et al., 2022; Sutton & DeRose, 2021) and the immunofluorescence experiments described above, we chose a 3 h treatment time point for our ChIP-seq experiments as this represented an early timepoint where nucleolar stress and inhibition of rRNA synthesis were observed in oxaliplatin treatments.

ChIP-seq analysis of Pol I (RPA194) in untreated A549 cells revealed broad enrichment of Pol I throughout the promoter and transcribed regions of rDNA, which sharply decreased in the intergenic spacer (IGS) (**Figure 3A**). ChIP-seq maps of Pol I-rDNA occupancy in BMH-21 and oxaliplatin treatments displayed a general loss of occupancy in A549 cells, consistent with inhibition of Pol I activity and loss of rRNA synthesis. Both oxaliplatin and BMH-21 treatments induced Pol I disengagement from rDNA in A549 cells, with a relative depletion in Pol I occupancy of ∼50% for oxaliplatin, and ∼70% for BMH-21 (**Figure 3B**). The broad depletion in Pol I occupancy following oxaliplatin treatments suggests the loss of 47S pre-rRNA transcripts previously measured by metabolic labeling (Pigg et al., 2022) result from Pol I inhibition. The dramatic reduction in Pol I occupancy by BMH-21 is likely accompanied by BMH-21 induced degradation of RPA194 (Jacobs, Huffines, et al., 2022). Cisplatin, in contrast, induced a slight reduction in total relative Pol I occupancy.

**Figure 3.**
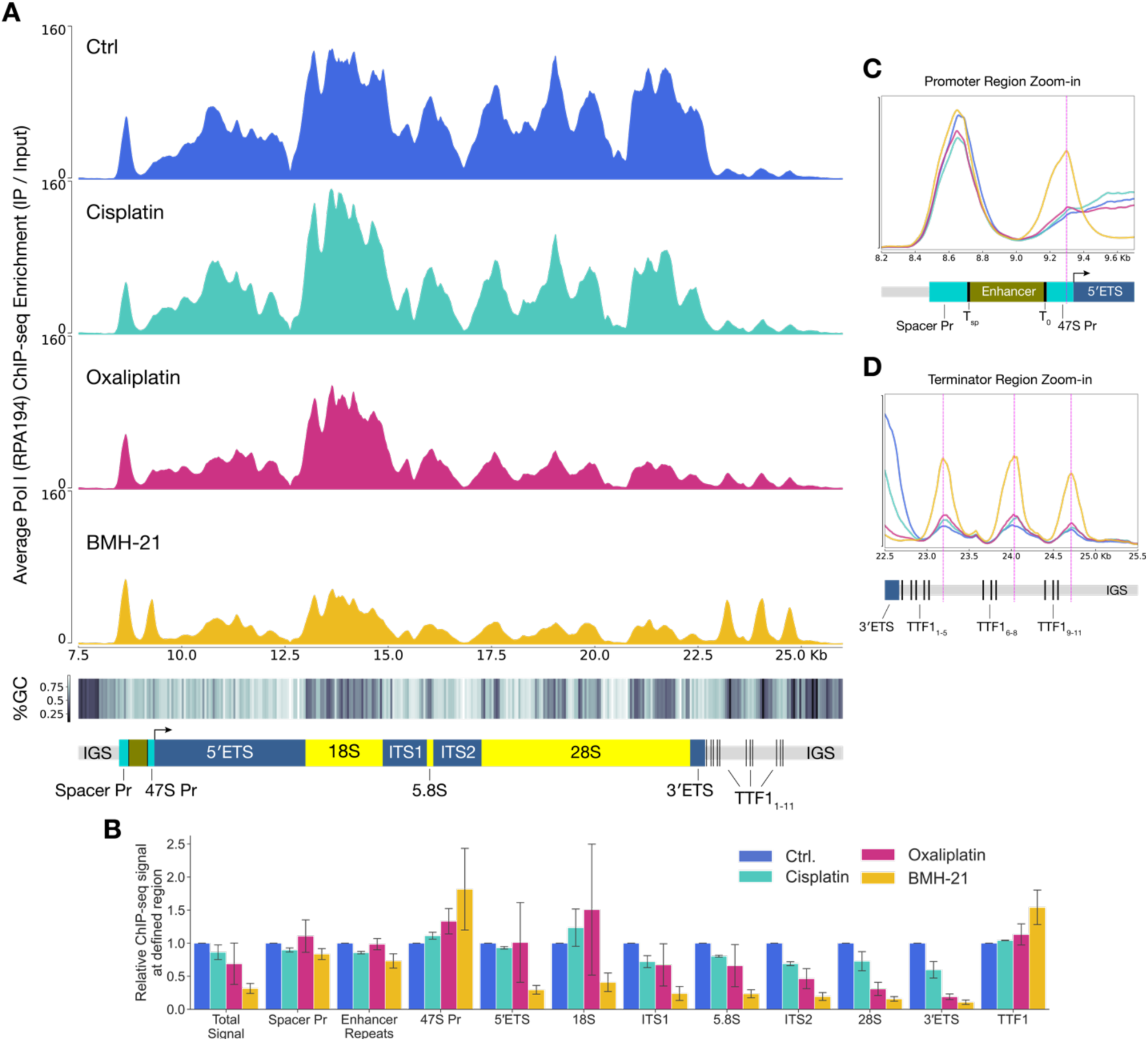
Platinum chemotherapeutics and BMH-21 treatment influence rDNA occupancy of Pol I. **(A)** ChIP-seq mapping of Pol I (RPA194) occupancy along the rDNA gene in A549 cells, following 3 h treatment with 10 µM cisplatin or oxaliplatin, or 1 µM BMH-21. Average sequence coverage is normalized to the read count per million reads (CPM) and shown as enrichment over the input for the associated treatment. Percent GC-content (%GC) is calculated from a 50-bp sliding window at each bp across the length of the rDNA gene. Intergenic spacer region (IGS), Spacer and 47S promoter (Spacer Pr, 47S Pr), 18S and 28S rRNA genes, internal transcribed spacers (ITS1, ITS2), transcription termination factor binding sites (TTF1_1-11_), and external transcribed spacers (5′ETS and 3′ETS), are labeled on the rDNA gene shown below, arrow indicates transcription start site. **(B)** Relative Pol I ChIP-seq signal mapped at defined regions of rDNA. Total read density within each defined rDNA region is normalized to the ChIP-seq signal in untreated cells. Plotted as mean ± SEM, n = 2 replicates. Zoom-in of Pol I rDNA occupancy in the **(C)** promoter region and **(D)** terminator region of rDNA shown in **(A)**. Enhancer Repeats (Enhancer), TTF1 initial binding sites (T_sp_ and T_0_), and termination binding sites (TTF1_1-11_) are labeled below.

Pol I occupancy maps in U2OS cells showed similar decrease in occupancy following BMH-21 treatments compared to A549 cells, (**Figure S1**) leading to a ∼60% reduction in relative Pol I occupancy (**Figure S1**). However, treatment with oxaliplatin in U2OS cells did not cause any significant change in Pol I occupancy, and cisplatin treatments lead to a nearly 2-fold enrichment in Pol I occupancy in comparison with untreated cells (**Figure S1**).

ChIP-seq occupancy mapping of oxaliplatin-treated A549 cells did not reveal a distinct pause site or enrichment motif, suggesting that Pol I inhibition does not result from a specific molecular disruption of the Pol I complex-rDNA interaction. However, oxaliplatin treatment did induced a slight decreasing gradient in relative Pol I occupancy in the 5′ to 3′ direction (**Figure 3A**) which was also observed in BMH-21 occupancy measurements, and previously reported with ActD (Jean-Clément Mars et al., 2020).

ChIP-seq analysis of UBF in untreated A549 and U2OS cells showed expected rDNA enrichment patterns throughout the promoter and transcribed regions of rDNA, which sharply decreased in the IGS (**Figure 4A** and **S2**). Despite the strong correlation between UBF occupancy and the rate of rRNA synthesis (Sanij et al., 2008; Theophanous et al., 2023), we did not observe a strong decrease in UBF occupancy in A549 and U2OS cells following treatment with oxaliplatin or BMH-21, and UBF occupancy patterns or specific enrichment sites did not appear to be significantly affected by treatment with any compound.

**Figure 4.**
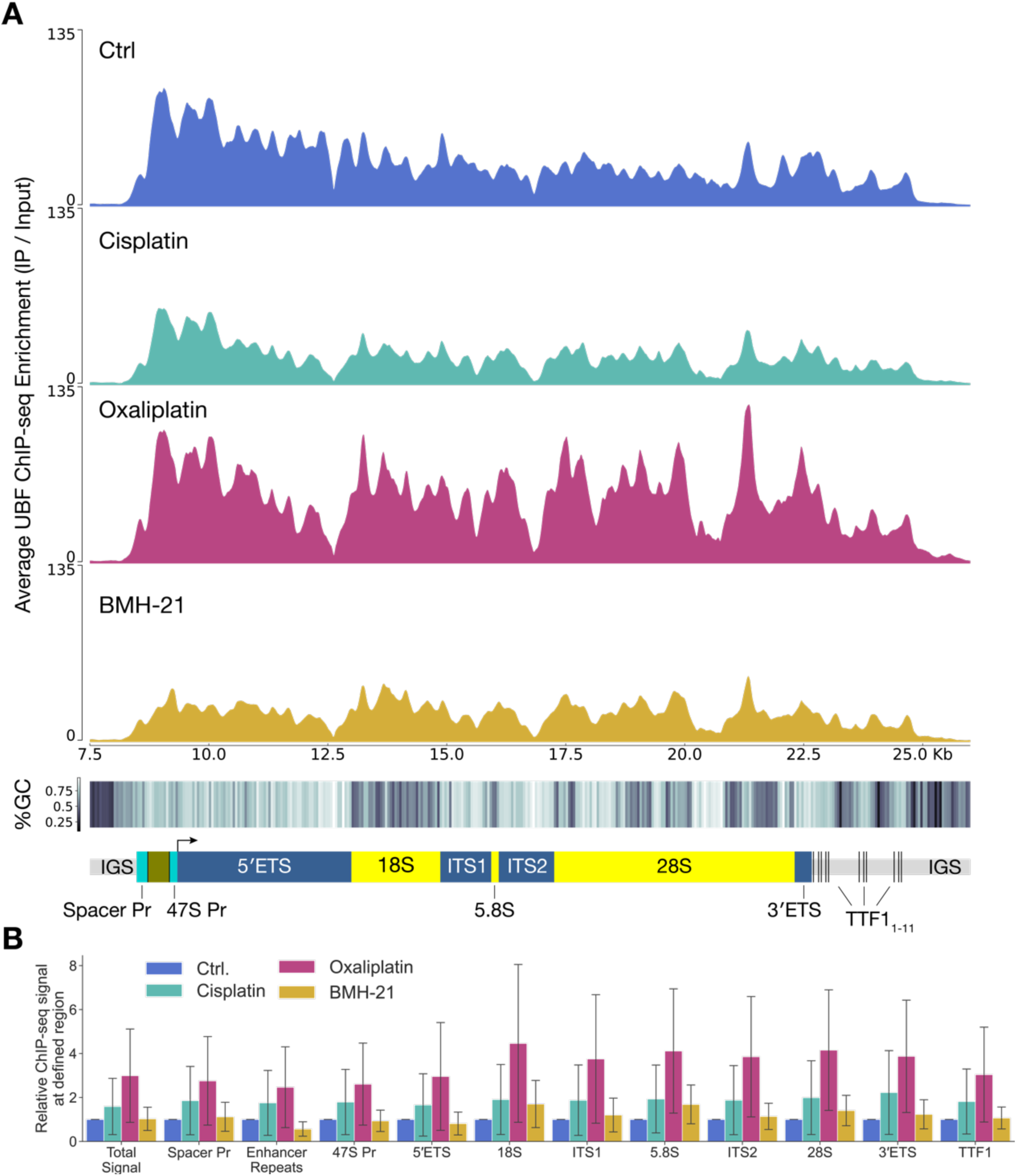
Influence of platinum chemotherapeutics and BMH-21 treatment on UBF-rDNA occupancy. **(A)** ChIP-seq mapping of UBF occupancy along the rDNA gene in A549 cells, following 3 h treatment with 10 µM cisplatin or oxaliplatin, or 1 µM BMH-21. Average sequence coverage is normalized to the read count per million reads (CPM) and shown as enrichment over the input sequence for the associated treatment. Diagram of rDNA gene aligned below: intergenic spacer region (IGS), spacer and 47S promoter (Spacer Pr, 47SPr), internal transcribed spacers (ITS1, ITS2), transcription termination factor biding sites (TTF1_1-11_), external transcribed spacers (5′ETS and 3′ETS), arrow indicates transcription start site. **(B)** Relative Pol I ChIP-seq signal mapped at defined regions of rDNA. Total read density within each defined rDNA region is normalized to the ChIP-seq signal in untreated cells. Plotted as mean ± SEM, n = 2 replicates.

In A549 cells, relative UBF occupancy increased substantially in oxaliplatin treatments, showing ∼1.5× overall enrichment compared to untreated cells (**Figure 4B**). BMH-21 and cisplatin induced a moderate 30% depletion in UBF occupancy. In contrast, cisplatin and oxaliplatin treatments in U2OS cells revealed a substantial increase in overall rDNA enrichment, ∼2× and 1.8× respectively, when compared to untreated cells, whereas BMH-21 treatments induced a slight 20% reduction in relative UBF occupancy (**Figure S2**).

Enrichment in UBF occupancy by cisplatin and oxaliplatin treatments may be partially due an unusually high affinity for Pt-DNA adducts, which has been reported for UBF (Hamdane et al., 2015; Zhai et al., 1998). More recent studies suggest UBF binding to rDNA is unperturbed by cisplatin or oxaliplatin treatments (Nechay et al., 2023).

However, the lack of specific UBF enrichment sites along the rDNA gene or correlation with G-rich regions (see below) suggests the increase in UBF occupancy observed in cisplatin and oxaliplatin treatment may be influenced by more global impacts on rRNA transcription and rather than recognition of specific rDNA-adducts.

### BMH-21 induces Pol I stalling along rDNA

Overall Pol I occupancy decreased under treatment with BMH-21, however BMH-21 treatment also led to a relative enrichment in Pol I occupancy at the 47S promoter region and downstream of the TTF1 binding clusters (**Figure 3C** and **D**). The relative occupancy of Pol I within the 47S promoter region increased approximately 2-fold following BMH-21 treatment in A549 cells (**Figure 3B**), which peaked approximately 50 bp upstream of the transcription start site or 125 bp downstream of the TTF1 binding site, T_0_ (**Figure 3C**). These enrichment peaks may indicate a specific block or stalling of Pol I bound to rDNA. Stalling of the Pol I initiation complex aligns with early findings in *S. cerevisiae* that BMH-21 primarily disrupts transcription initiation and early elongation phase (Jacobs, Fuller, et al., 2022; Jacobs, Huffines, et al., 2022), and may demonstrate a specific mode for Pol I inhibition induced by BMH-21 at pre-initiation or early elongation steps. Additional occupancy mapping of Pol I transcription factors involved in polymerase initiation, such as the Pol I-specific transcription initiation factor, RRN3 or selective factor 1 (SL-1) may elucidate the specific step of inhibition.

Relative enrichment in Pol I occupancy at TTF1 binding sites, TTF1_1-11_ was also observed in BMH-21 treated A549 cells (**Figure 3D**). TTF1 is an rDNA specific transcription termination factor which binds to Sal Box terminator elements downstream of the 3′ end of the pre-rRNA. TTF1 binding induces pausing of the Pol I complex, which mediates RNA Pol I transcription termination by promoting polymerase release (Németh et al., 2008). Peaks in RPA194 occupancy in the TTF1 region were observed approximately 60–80 bp downstream of TTF1 binding clusters, which suggests that Pol I termination, release of the termination complex, or both may also be disrupted by BMH-21. To our knowledge, BMH-21 induced disruptions in Pol I transcription termination or polymerase release have not been otherwise reported, and may represent an additional mode of late stage disruption of Pol I which warrants further study.

In contrast, for oxaliplatin treatments, the lack of a specific enrichment pattern in Pol I-rDNA occupancy during early stages of nucleolar stress and loss of rRNA transcription suggest that Pol I inhibition may not be primarily caused by a molecular disruption of Pol I interaction with a specific region of rDNA. The loss of rDNA occupancy may reflect an early nucleolar stress response leading to a down regulation of rRNA synthesis. Alternatively, oxaliplatin may induce a more specific perturbation of Pol I function at earlier timepoints, prior to the earlier stages of nucleolar stress, which may lead to the downstream inhibition of rRNA transcription and general loss of Pol I occupancy across the rDNA gene as observed here.

### Pol I and UBF occupancy measured by ChIP-seq do not correlate to relative rDNA GC-content

BMH-21 and other specific Pol I inhibitors are thought to gain specificity through their preferential interaction with GC-rich regions of the genome, such as those found in the coding regions of rDNA (Peltonen et al., 2014; Pitts & Laiho, 2022). BMH-21 has previously been demonstrated though NET-seq in *S. cerevisiae* to induce Pol I stalling upstream of G base pairs in rDNA coding region, (Jacobs, Huffines, et al., 2022) consistent with the hypothesis that GC-rich sequences direct Pol I specificity in that small molecule inhibitor.

In our ChIP-seq mapping data, we were unable to find a correlation between enrichment in Pol I or UBF occupancy and relative GC-content within the rDNA (**Figures 3A and 4A**) and found no evidence for a sequence-specific buildup of occupancies that would suggest specific disruptions in Pol I engagement which could result in perturbed rRNA transcription, at the resolution of these experiments. However, ChIP-seq rDNA occupancy mapping may be insufficient to detect these small nucleotide sequence motifs, in comparison to higher resolution NET-seq experiments, due to the long read lengths required for ChIP-seq occupancy mapping and potential biases in sequencing GC-rich fragments. These factors make it difficult to conclude from these ChIP-seq experiments whether GC-content plays a meaningful role in directing Pol I or UBF specificity.

In a broader context, our ChIP-seq experiments also underscore the high degree of sequence specificity of Pol I transcription factors for the transcribed regions of rDNA, which is maintained despite the dramatic relocalization events which take place during nucleolar stress. These protein–nucleic acid interactions may be an important factor in further understanding the complex interactions between biomolecules which maintain nucleolar homeostasis and direct response to cellular stress.

## Methods

### Reagents and tools

| Reagent | Source | Cat. |
| --- | --- | --- |
| Cisplatin | AvaChem Scientific | 1714 |
| Oxaliplatin | TCI Chemicals | O0372 |
| BMH-21 | Sigma-Aldrich | SML1183 |
| Mouse monoclonal anti-UBF (F-9) | Santa Cruz Biotechnology | sc-13125 |
| Mouse monoclonal anti-RPA194 (C-1) | Santa Cruz Biotechnology | sc-48385 |
| Mouse IgG Isotype Control | Invitrogen | 31903 |
| Goat anti-Mouse Alexa Fluor 488 | Abcam | Ab150133 |

### Cell culture and Treatment

A549 human lung carcinoma cells (ATCC, #CCL-185) were cultured at 37°C, 5% CO_2_ in Dulbecco’s Modified Eagle Medium (DMEM) supplemented with 10% Fetal Bovine Serum (FBS) and 1% antibiotic-antimycotic. U2OS human bone osteosarcoma cells (ATCC, #HTB-96) were cultured at 37°C, 5% CO_2_ in McCoy’s 5A medium supplemented with 10% Fetal Bovine Serum (FBS) and 1% antibiotic-antimycotic.

Stock solutions of cisplatin (5mM, 0.9% NaCl) and oxaliplatin (5mM, ddH_2_O) were prepared fresh on the day of treatment. BMH-21 stock solution (500 µM, DMSO) was stored in aliquots at –20°C and thawed prior to use. Stock solutions were diluted into pre-warmed cell media immediately prior to treatment. All drug treatments were conducted on cells between passages 8–20 at 80% confluency and carried out at a final concentration of 10 µM for cisplatin and oxaliplatin, and 1 µM for BMH-21.

### Immunofluorescence

Cells were grown on #1.5, 10 mm dia. coverslips (Ted Pella no. 260368) in a 24-well plate as described above. Following treatment, cells were washed twice with phosphate buffered saline (PBS) then fixed for 20 minutes at RT with 4% paraformaldehyde (PFA) in PBS and permeabilized with 0.5% Triton-X in PBS for 20 min at RT. Coverslips were then blocked with 1% bovine serum albumin (BSA) in PBST (PBS with 0.1% Tween-20) for 10 min, twice and incubated with primary antibody for 1 h. Coverslips were washed 3x with PBS for 5 min each at RT, and then incubated with secondary antibody (1:1000, in PBST with 1% BSA) for 1 h in the dark. Following three more washes with PBS for 5 min each, coverslips were mounted on slides with ProLong Diamond Antifade Mountant with DAPI (ThermoFisher, cat no. P36971) and allowed cured at RT overnight before imaging. Primary antibodies and concentrations: anti-RPA194, (1:500, in PBST with 1% BSA) and anti-UBF (1:400, in PBST with 1% BSA)

### Chromatin Immunoprecipitation

Chromatin Immunoprecipitation was performed according to (Sullivan & Santos, 2020) with some modification. Following treatment, cells were washed twice with cold PBS, harvested with 2 mL of TrypLE (Gibco, cat no. 12604013), and collected in 6 mL of cell media. Cells were crosslinked with 500 µL of 16% PFA (final concentration, 1% PFA) for 10 min at RT with rotation, which was quenched with 850 µL of cold 1.25M glycine (final concentration, 125 mM glycine) for 5 min at RT with rotation.

Crosslinked cells were pelleted (800 × *g*, 6 min, 4°C) and washed twice with cold PBS, then resuspended in 1 mL PBS for cell counting using a TC20 Automated Cell Counter (Bio-Rad), at 1:1 dilution with trypan blue. Approx. 5 × 10^6^ cells were transferred to a 1.5mL protein lo-bind tube, pelleted (800 × *g*, 6 min, 4°C), and then resuspended in 200 µL of cold cell lysis buffer (800mM NaCl, 25 mM Tris pH 7.5, 5mM EDTA, 1% Triton X-100, 0.1% SDS, and 0.5% sodium deoxycholate) supplemented with protease/phosphatase inhibitor (Cell Signaling Technology, #5872S). Cells snap-frozen in dry ice and acetone and stored at –80°C.

On the day of experiment, cell lysate was thawed on ice and sheared using a Bioruptor sonicator (Diagenode) at 4°C for 20 cycles of 30 sec ON/OFF. Sheared chromatin was diluted with 200 µL of cold chromatin dilution buffer (25 mM Tris pH 7.5, 5mM EDTA, 1% *v/v* Triton X-100, 0.1% *w/v* SDS) and clarified by centrifugation (13,600 × *g*, 30 min, 4°C). A 20 µL of diluted chromatin (5% Input) was aliquoted and stored at – 20°C to serve as the ChIP Input. For each IP reaction, 100 µL of diluted chromatin was combined with 50µL of preincubated Dynabeads–antibody mix in a 1.5 mL protein lo-bind tube and incubated overnight at 4°C with rotation. Dynabeads–antibody mix was prepared as followed: For each IP, 25 µL of Dynabeads Protein G magnetic beads (ThermoFisher, #1003D) was combined with 12.5 µL of ChIP antibody (anti-RPA194 or anti-UBF) or 1.875 µL of IgG control, and diluted chromatin dilution buffer to a final volume of 50 µL, which was incubated at RT for 2 h with rotation.

The following day, chromatin-bead mixture was set on a magnetic stand and the beads were then washed at 4°C with 500 µL of IP wash buffer—rotating for 5 min in between each of the following wash steps: *i*) “low salt” buffer (140 mM NaCl, 50 mM HEPES pH 7.9, 1mM EDTA, 1% Triton X-100, 0.1%SDS, 0.1% sodium deoxycholate) *ii*) “high salt” buffer (500 mM NaCl, 50 mM HEPES pH 7.9, 1mM EDTA, 1% Triton X-100, 0.1%SDS, 0.1% sodium deoxycholate) *iii*) LiCl buffer (20 mM Tris pH 7.5, 1mM EDTA, 250mM LiCl, 0.5% Triton X-100, 0.5% sodium deoxycholate) and *iv*) TE buffer (10 mM Tris pH 7.5, 1mM EDTA). After the final wash step the beads were incubated with 100 µL of elution buffer (10 mM Tris pH 7.5, 1mM EDTA, 1% SDS) at 65°C for 5 min on thermo shaker (VWR), followed by an additional 15 min at RT. The bead mixture was set on a magnetic stand and supernant was collected a new centrifuge tube. The elution step was repeated to obtain 200 µL of ChIP chromatin.

To reverse crosslinking and digest contaminates, input samples were first thawed on ice and diluted with 180 µL with chromatin dilution buffer. 8 µL of 4 M NaCl (final concentration, 160 mM) and 0.5 µL of RNase A (ThermoFisher, #EN0531) (final concentration, 23 µg/mL) were added to both ChIP and Input samples, followed by incubation at 65°C for 8 h in a thermocycler (Eppendorf). Next, 2 µL of 0.5 M EDTA (final concentration, 5 mM) and 2 µL Proteinase K (New England Biolabs, #P8107S) (final concentration, 200 µg/mL) were added to each sample and incubated in a thermocycler 45°C for 2 h. ChIP and Input DNA was purified using a Zymo ChIP DNA Clean & Concentrator Kit (#D5205) and eluted into 25 µL of DNA Elution buffer (10 mM Tris pH 8.5, 0.1 mM EDTA).

DNA library for next-gen sequencing was prepared using the NEBNext Ultra II DNA Library Prep Kit, for Illumina (New England Biolabs, #E7600S), multiplexed using NEBNext Ultra II DNA Library Prep Kit, for Illumina (New England Biolabs, #E7645S) and sequenced on an Illumina NovaSeq 6000 platform.

### Data processing and rDNA mapping

A custom human genome assemble for rDNA mapping (*hs1-rDNA_genome_v1.0*), along with annotation files were obtained from publicly available GitHub repository (https://github.com/vikramparalkar/rDNA-Mapping-Genomes) based on published methods (George et al., 2023). The *hs1* (T2T-CHM13) reference human genome was masked for rDNA-matching loci, and a single full-length human rDNA sequence, KY962518.1 (44,838 nt) (Kim et al., 2018) was then added as an extra chromosome (chrR).

Demultiplexed FASTQ read files were first trimmed with Trimmomatic (v0.39) (Bolger et al., 2014) with the following parameters: ILLUMINACLIP:TruSeq3-PE.fa:2:30:10:2:True LEADING:3 TRAILING:3 SLIDINGWINDOW:4:15 MINLEN:30, then mapped with Bowtie2 (v2.5.4) (Langmead & Salzberg, 2012) with the following parameters: –X 2000 –k 3 to allow for indel variations up to 2 kb, documented in rDNA repeats (Jean-Clement Mars et al., 2018), and to permit for multiple alignments for each sequence read. The resulting SAM files were filtered and into the BAM format using samtools (v1.19) (Li et al., 2009) with parameters: – F 4 –q 1 to retain multi-mapped read, then sorted and indexed with samtools sort and index.

Normalized coverage tracks were generated from indexed BAM files with deeptools (v3.5.3) (Ramírez et al., 2016) bamCoverage with the following parameters: –bs 1 –-smoothLength 15 –-maxFragmentLength 500 –-minFragmentLength 30 –-centerReads –-extendReads –-normalizeUsing CPM –-ignoreForNormalization chrX chrM. Reads were normalized as total counts per million (CPM) (*# of reads per bin / total reads*) to account for differences in sequencing depth.

Averaged coverage tracks were obtained using deeptools (v3.5.3) (Ramírez et al., 2016) bigWigAverage with the following parameters: –bs 1 –r chrR. “IP Enrichment” was calculated using deeptools (v3.5.3) (Ramírez et al., 2016) bigwigCompare from average coverage tracks as the ratio of ChIP reads to Input DNA reads at each base position (*ChIP coverage* / *Input coverage*). Coverage track figures were generated with pyGenomeTracks (Lopez-Delisle et al., 2021).

### Data Availability

Raw and processed ChIP-seq data from this publication have been deposited to the NCBI Gene Expression Omnibus (GEO; https://www.ncbi.nlm.nih.gov/geo/) under the accession number GSE284654.

## Acknowledgements

This investigation was supported by a grant from the National Science Foundation (CHE 2109255) to V.J.D. and a National Institutes of Health Training Grant to M.V.Y. (T32 GM007759), and by the Genomics & Cell Characterization Core Facility at the University of Oregon.

