## Supplemental Information for "Nucleolar Stress-inducing Compounds Influence rDNA occupancy of RNA Polymerase I Transcription Machinery"

**Figure S1.** Influence of platinum chemotherapeutics and BMH-21 treatment on rDNA occupancy of Pol I in U2OS cells

**Figure S2.** Influence of platinum and BMH-21 treatment on UBF-rDNA occupancy in U2OS cells

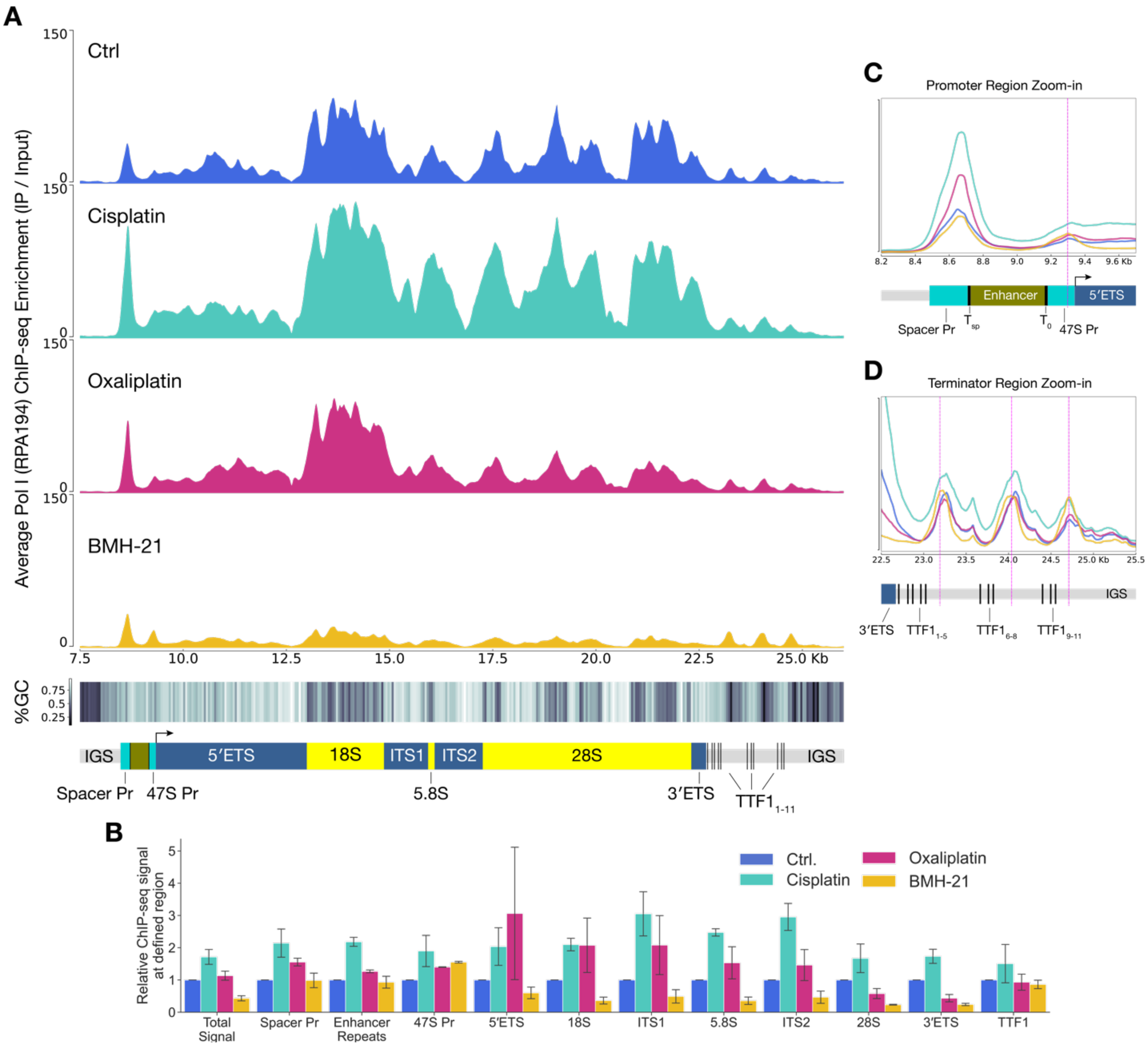


**Figure S1.** Influence of platinum chemotherapeutics and BMH-21 treatment on rDNA occupancy of Pol I. **(A)** ChIP-seq mapping of Pol I (RPA194) occupancy along rDNA gene in U2OS cells, following 3 h treatment with 10 $\mu$M cisplatin or oxaliplatin, or 1 $\mu$M BMH-21. Average sequence coverage is normalized to the read count per million reads (CPM) and shown as enrichment over the input sequence for the associated treatment. Percent GC-content (%GC) is calculated from a 50-bp sliding window at each bp across the length of the rDNA gene. Intergenic spacer region (IGS), Spacer and 47S promoter (Spacer Pr, 47S Pr), 18S and 28S rRNA genes, internal transcribed spacers (ITS1, ITS2), transcription termination factor binding sites ($TTF1_{1-11}$), and external transcribed spacers (5′ETS and 3′ETS), are labeled on the rDNA gene shown below, arrow indicates transcription start site. **(B)** Relative Pol I ChIP-seq signal mapped at defined regions of rDNA. Total read density within each defined rDNA region is normalized to the ChIP-seq signal in untreated cells. Plotted as mean ± SD, n = 2 replicates. Zoom-in of Pol I rDNA occupancy in the **(C)** promoter region and **(D)** terminator region of rDNA shown in **(A)**. Enhancer Repeats (Enhancer),TTF1 space promoter and initial binding sites ($T_{sp}$ and $T_0$) and termination binding sites ($TTF1_{1-11}$) are labeled below.

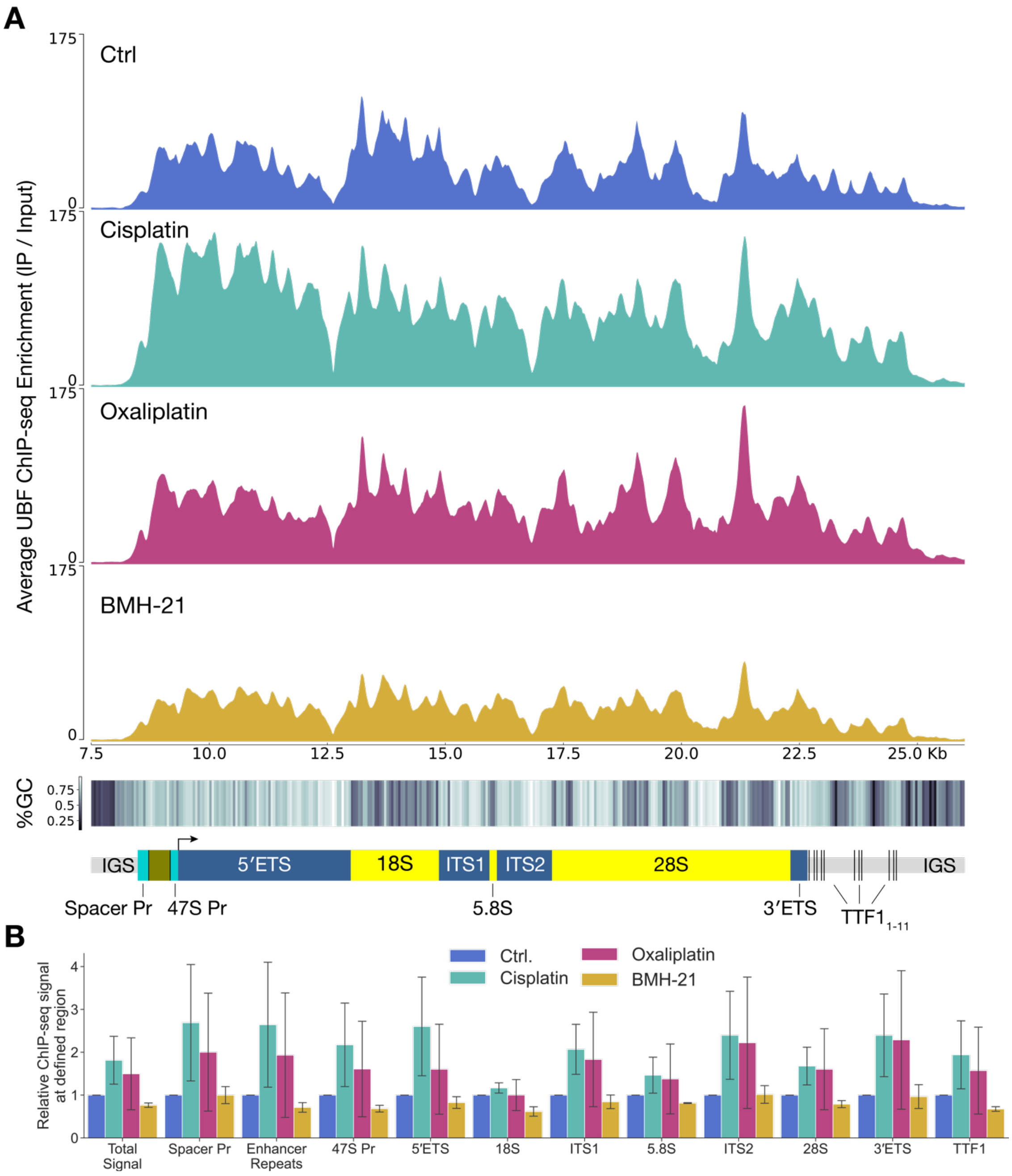


**Figure S2.** Influence of platinum and BMH-21 treatment on UBF-rDNA occupancy. **(A)** ChIP-seq mapping of UBF-rDNA occupancy in U2OS cells, following 3 h treatment with 10 $\mu$M cisplatin or oxaliplatin, or 1 $\mu$M BMH-21. Average sequence coverage is normalized to the read count per million reads (CPM) and shown as enrichment over the input sequence for the associated treatment. Percent GC-content (%GC) is calculated from a 50-bp sliding window at each bp across the length of the rDNA gene. Diagram of rDNA gene aligned below: intergenic spacer region (IGS), spacer and 47S promoter (SpPr, 47SPr), internal transcribed spacers (ITS1, ITS2), transcription termination factor biding sites ($TTF1_{1-11}$), external transcribed spacers (5′ETS and 3′ETS), arrow indicates transcription start site. **(B)** Relative Pol I ChIP-seq signal mapped at defined regions of rDNA. Total read density within each defined rDNA region is normalized to the ChIP-seq signal in untreated cells. Plotted as mean ± SD, n = 2 replicates.
